# Systemically delivered *Bacteroides thetaiotaomicron*-derived bacterial extracellular vesicles inhibit primary and metastatic melanoma growth

**DOI:** 10.64898/2026.05.20.726595

**Authors:** Christopher A. Price, Emily J. Jones, Nilda Ilker, Alicia Nicklin, Rokas Juodeikis, Alastair M. McKee, Luke Mitchell, Régis Stentz, Lindsay J. Hall, Simon R. Carding, Stephen D. Robinson

## Abstract

The gut microbiome can contribute to anti-tumour immunity and cancer therapy responses, but translating live microbe-based interventions remains challenging due to safety, controllability, and delivery constraints. Bacterial extracellular vesicles (BEVs) are an attractive cell-free alternative, as they package bacterial cargo into a nanoscale format capable of host-cell engagement, immunological activation, and systemic distribution. Here, we investigated the anti-tumour potential of BEVs derived from the human gut commensal *Bacteroides thetaiotaomicron* (Bt). We show that delivery route is a major determinant of efficacy. Intravenous, but not intraperitoneal, administration produced robust anti-tumour activity in a B16F10 melanoma mouse model. Intravenously delivered Bt BEVs suppressed primary tumour growth in a dose-dependent manner and reduced metastatic outgrowth in the lung. Bt BEVs did not directly impair tumour-cell viability *in vitro*, but they activated NF-κB and Toll-like receptor signalling in innate immune reporter systems and localised to tumour tissue following systemic administration. Together, these data support a model in which Bt BEVs act via host immune modulation rather than direct tumour cytotoxicity. These findings identify naturally produced commensal-derived Bt BEVs as a potential microbial therapeutic modality and as an alternative to the use of live bacterial administration in cancer therapy.

## Introduction

The intestinal microbiome is emerging as an important modifier of cancer progression and treatment response, with evidence that microbial communities and their products can shape anti-tumour immunity, alter treatment efficacy, and influence toxicity(*1-3*). Emphasis is now shifting from descriptive taxonomic associations to the identification of defined microbial effectors with demonstrable therapeutic properties(*4, 5*).

One candidate class of such effectors is bacterial extracellular vesicles (BEVs), which are naturally produced by both Gram-negative and Gram-positive bacteria (*6, 7*). Bacterial extracellular vesicles carry a range of bacterial cargo, including membrane-associated and cytosolic components, and can interact with epithelial and immune cells at mucosal surfaces in the gastrointestinal tract (GIT)(*8-10*). This provides a plausible cell-free mechanism by which bacterial products can influence host responses without requiring direct bacterial colonisation or invasion. Via the GIT, BEVs can also gain access to the circulatory system to reach distant tissues and organ systems, including the brain(*11*). In oncology, BEVs have increasingly been explored as immunostimulatory agents, tumour-targeting carriers, and vaccine or drug-delivery platforms(*6, 8, 12*). However, much of this literature has focused on engineered bacteria and vesicles, or vesicles derived from pathogens, rather than native vesicles released by common commensals. As a result, the anti-tumour potential of naturally produced vesicles from non-pathogenic members of the human microbiome remains less well defined.

*Bacteroides thetaiotaomicron* (Bt) is a prominent human gut commensal Gram-negative bacterium with well-established effects on host physiology and mucosal immune homeostasis(*13-15*). Bt BEVs are naturally produced in the GIT, can cross the intact intestinal epithelial barrier(*11*), and can modulate immune(*16*), epithelial(*17*), and neuronal cell(*18*) responses. This supports the idea that biologically relevant host effects can be mediated by secreted non-viable microvesicles rather than by live bacterial colonisation alone(*16, 17*). Thus, Bt is a credible source organism for testing whether native commensal bacteria-derived BEVs can function as non-viable anti-cancer therapeutics.

Despite their promise, several key questions remain unresolved regarding BEVs. It is unclear whether native Bt BEVs can generate anti-tumour effects *in vivo*, whether the route of administration influences efficacy, what their mechanism of action might be, whether direct tumour toxicity or host immune modulation are mechanistically involved. These questions are relevant for clinical translation, as formulation and delivery parameters will be major determinants of success for vesicle-based microbial therapeutics(*19, 20*).

Here, we evaluated the anti-tumour activity of Bt-derived BEVs in preclinical tumour models, focusing on delivery route and *in vivo* efficacy. We show that intravenous (IV), but not intraperitoneal (IP), administration produces dose-dependent suppression of primary tumour growth and reduced pulmonary lesion outgrowth. We further show that Bt BEVs activate innate immune signalling pathways *in vitro* and localise to tumour tissue after systemic delivery. Together, these findings position native commensal Bt BEVs as a potential microbial therapeutic modality, identifying delivery route as a critical determinant of biological activity.

## Methods

### Animal studies

All animal procedures were performed in accordance with UK Home Office regulations and the European Legal Framework for the Protection of Animals used for Scientific Purposes (European Directive 86/609/EEC). C57BL/6 mice were maintained at the Disease Modelling Unit, University of East Anglia, under project licence PP8873233. Mice were used at 8–12 weeks of age and were randomly mixed between cages before the start of experiments. Subcutaneous tumour experiments were performed in age-matched male C57BL/6 mice, whereas the B16F10 experimental metastasis model used age-matched male and female C57BL/6 mice.

### Bacterial culture, BEV generation, and quantification

*Bacteroides thetaiotaomicron* VPI-5482 was grown anaerobically at 37°C in Bacteroides Defined Media (BDM) adapted from *Juodeikis et al*.(*21*). BEVs were isolated from culture supernatants as previously described by *Stentz et al*.(*22*) with minor modifications. Briefly, bacterial cultures were centrifuged at 5,500 × g for 45 min at 4°C, and the supernatants were passed through 0.22 μm polyethersulfone membranes (Sartorius). Filtered supernatants were concentrated by ultrafiltration using a 100 kDa molecular weight cut-off device (Vivaflow 50R, Sartorius), washed with 500 mL PBS, concentrated to 1 mL in sterile PBS, and passed through a final 0.22 μm syringe filter (Sartorius). BEV preparations were stored at 4°C, and sterility was confirmed by plating on BHI–hemin agar. Particle size and concentration were determined by nanoparticle tracking analysis using the ZetaView platform (ZetaView, Particle Matrix).

### Nanoparticle tracking analysis (ZetaView)

The size distribution and concentration of bacterial extracellular vesicles were determined by nanoparticle tracking analysis using a ZetaView PMX 220 TWIN instrument (Particle Metrix, Germany) operated according to the manufacturer’s instructions. Vesicle suspensions were diluted up to 50,000-fold in particle-free water to obtain approximately 50–200 particles per frame. For each sample, videos were acquired at 11 positions with two measurement cycles per position (60 frames per position; 2 s video length; temperature 25°C; camera sensitivity 80; shutter 100). Data were analysed using ZetaView NTA software version 8.05.12, with post-acquisition settings of minimum brightness 20, minimum area 5, maximum area 2,000, and trace length 30.

### Cell lines and culture conditions

B16F10 melanoma and CMT19T lung carcinoma cells were maintained in high-glucose DMEM supplemented with 10% fetal bovine serum and 100 U/mL penicillin/streptomycin. Cells were cultured at 37°C in 5% CO_2_ in flasks coated with 0.1% porcine gelatin.

The human myeloid leukaemia cell line THP-1-Blue (Invivogen) expressing an NF-κB-inducible secreted alkaline phosphatase (SEAP) reporter was maintained in RPMI-1640 supplemented with 10% fetal bovine serum, 1% penicillin/streptomycin, 100 μg/mL Normocin, and 10 μg/mL blasticidin. The human embryonic kidney cell line 293 (HEK293) expressing human TLR2 and hTLR4 inducible SEAP reporter (HEK-Blue) was maintained in RPMI-1640 supplemented with 10% fetal bovine serum, 1% penicillin/streptomycin, 100 μg/mL Normocin, and 1× HEK-Blue selection cocktail.

### THP1-Blue NF-κB reporter assay

THP1-Blue cells were seeded at 1 × 10^5^ cells per well in 96-well plates and incubated for 24 h with BEVs or LPS as a positive control. NF-κB activity was quantified using QUANTI-Blue detection medium according to the manufacturer’s instructions, and absorbance was measured at 650 nm.

### HEK-Blue TLR2 and TLR4 reporter assays

HEK-Blue hTLR2 and HEK-Blue hTLR4 cells were seeded at 5 × 10^4^ cells per well in 96-well plates in HEK-Blue detection medium containing BEVs or control ligands. LTA-BS (from *B. subtilis*; Invivogen) and LPS (from *E. coli* O111:B4; Sigma) were used as positive controls for TLR2 and TLR4, respectively. Cells were incubated for 16h at 37°C in 5% CO_2_, and reporter activity was quantified by absorbance at 650 nm measured on a VERSAmax microplate reader (Molecular Devices, San Jose, USA).

### Tumour-cell viability assay

To assess whether Bt BEVs directly altered tumour-cell viability, B16F10 cells were seeded at 2,000 cells per well in 96-well plates and allowed to adhere for 24 h. Cells were then treated with a range of BEVs for a further 24 h. alamarBlue reagent was added for 4 h at 37°C in 5% CO_2_, and fluorescence intensity was measured at 590 nm on a VERSAmax microplate reader (Molecular Devices, San Jose, USA).

### Subcutaneous tumour models and BEV treatment

4 × 10^5^ B16F10 melanoma cells or 1 × 10^6^ CMT19T lung carcinoma cells were injected into the flank of 8–12-week-old male C57BL/6 mice in 100 μL PBS. Tumour volume was measured three times weekly by digital caliper from the onset of palpable growth and calculated as length × width^2^ × 0.52. B16F10 tumours were typically harvested on day 14 and CMT19T tumours on day 18.

For BEV treatment protocols, BEVs were administered either IP three times weekly or IV every 3 days from the onset of palpable tumour, as specified in the corresponding experimental design. For B16F10 route-comparison experiments, 1 × 10^10^ BEVs were delivered by either IP or IV injection. In IV dose-response studies, tumour-bearing mice received escalating BEV doses, with 1 × 10^10^ particles representing the highest and most effective dose tested. The same IV treatment schedule was used in the CMT19T model. Body weight was monitored throughout treatment.

### Experimental B16F10 metastasis model

1 × 10^6^ B16F10 cells were injected IV via the tail vein. Tumour cells were allowed to circulate and seed at metastatic sites before initiation of BEV therapy. BEV treatment began 3 days after tumour cell injection, and lungs were harvested on day 14 for quantification of metastatic nodule number and average nodule area.

### Tissue histology

To assess treatment-associated tissue toxicity, major organs were collected at endpoint and processed as formaldehyde-fixed paraffin-embedded tissue. Organs were fixed for 16 h in 4% paraformaldehyde at 4°C, dehydrated through graded ethanol, cleared in xylene, and embedded in paraffin. Paraffin blocks were sectioned at 6 μm using a rotary microtome, mounted onto positively charged glass slides, and incubated for 16 h at 37°C. Sections were then deparaffinised in xylene, rehydrated through decreasing concentrations of ethanol into water, and stained with haematoxylin and eosin using a Leica ST5020 tissue multi-stainer.Coverslips were mounted with Neo-Mount. Brightfield images were acquired using an Olympus BX60 microscope fitted with a Jenoptik C10 camera.

### BEV biodistribution

For *in vivo* tracking studies, NanoLuc-tagged Bt BEVs (BEV-NLuc) were generated by genetically inserting nanoluciferase into parental *Bacteroides thetaiotaomicron* VPI-5482 as previously described(*11, 23*), followed by BEV isolation as described above. For biodistribution experiments, 1 × 10^10^ BEV-NLuc or vehicle control was injected IV via the tail vein into day 10 B16F10 tumour-bearing mice. Mice were sacrificed 3 h post-dose, and organs were excised, immersed in sterile PBS, blotted dry, and arranged in large Petri dishes. Organs were then immersed in 10 mL furimazine substrate (Nano-Glo Luciferase Assay System, Promega) diluted 1:20 in PBS for 5 min before imaging on a Bruker In-Vivo Xtreme instrument using a 1 min bioluminescence exposure setting. Regions of interest were drawn around each organ, and bioluminescent signal was quantified in ImageJ.

### Statistical analysis

Data are presented as mean ± SEM unless otherwise indicated. Statistical analyses were performed using GraphPad Prism 9. Comparisons between two groups were performed using two-tailed unpaired t tests, with Welch’s correction applied where appropriate. Comparisons involving more than two groups were performed using ordinary one-way ANOVA with Dunnett’s multiple-comparisons test against the relevant vehicle control unless otherwise stated. P < 0.05 was considered statistically significant. Significance thresholds are indicated as *P < 0.05, ***P < 0.001 and ****P < 0.0001.

## Results

### Intravenous administration is required for robust anti-tumour activity of Bt BEVs

To define the vesicle preparation used for downstream functional testing, we first assessed the morphology and size distribution of Bt BEVs. Consistent with previous cryo-TEM characterisation of Bt BEVs generated using the same established preparation workflow, representative vesicles displayed rounded, membrane-bound morphology (Fig. 1A).Nanoparticle tracking analysis of two independent Bt BEV preparations generated on different days showed a number-weighted median diameter of 159 nm and mean particle diameter of 168 nm, with a distribution standard deviation of 67 nm (Fig. 1B). Together, these data support the recovery of nanoscale Bt BEV preparations suitable for *in vivo* functional testing.

**Figure 1.**
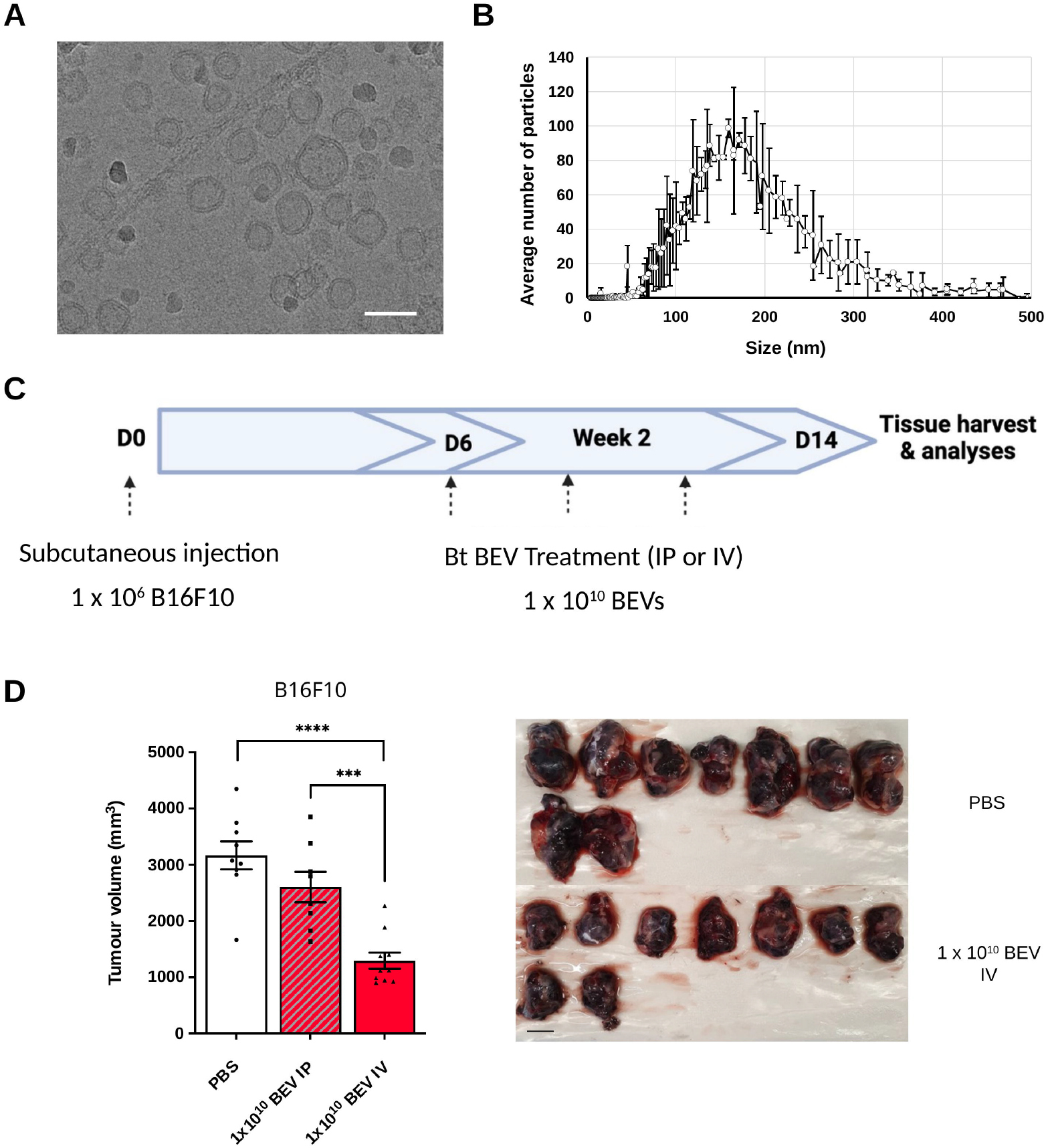
Bt BEVs are nanoscale membrane-bound vesicles and show route-dependent anti-tumour activity. (A)Representative cryo-transmission electron micrograph of Bt-derived BEVs generated using the established preparation workflow. Vesicles display rounded, membrane-bound morphology. Image adapted from *Juodeikis et al*. (2024)(*19*) (CC BY 4.0). Scale bar, 100 nm. (B)Nanoparticle tracking analysis (NTA) size distribution of Bt-derived BEVs. Particle size (nm) is plotted against the average number of particles per size bin for two independent samples prepared on different days. The combined number-weighted data gave a median (D50) size of 159 nm, with a mean of 168 nm and a standard deviation of 67 nm. (C) Experimental outline of the B16F10 primary melanoma route-comparison experiment. Mice were inoculated subcutaneously with 1 × 10^6^ B16F10 cells on day 0 and treated with 1 × 10^10^ Bt BEVs by either intraperitoneal (IP) or intravenous (IV) administration on experimental days 6, 9, and 11, followed by tissue harvest and analysis on day 14. (D) Quantification of endpoint B16F10 tumour volume following vehicle control, IP BEV, or IV BEV treatment, with representative endpoint tumour images shown at right. Scale bar = 1cm. Bars represent mean ± SEM, n = 8–10. Statistical significance was calculated by a two-tailed unpaired t-test. IP vs PBS, *P* = 0.1447; IV vs PBS, *P* < 0.0001; IV vs IP, *P* = 0.0004. \*\*\**P* < 0.001, \*\*\*\**P* < 0.0001.

To evaluate the therapeutic potential of Bt BEVs *in vivo*, we next assessed whether the delivery route influences anti-tumour efficacy. Bt BEVs were administered either IP or IV at an equivalent dose (1×10^10^ BEVs) in mice bearing established B16F10 melanoma tumours (Fig. 1C). While IP administration produced a non-significant trend reduction in tumour burden (*P* = 0.144), IV delivery resulted in a marked suppression of tumour growth relative to both PBS control (*P* < 0.0001) and IP-treated animals (*P* = 0.0004) (Fig. 1D).Quantification of endpoint tumour volumes demonstrated a substantially greater reduction following IV administration relative to IP treatment, indicating that delivery route is a key determinant of BEV efficacy. These findings establish that systemic administration is required to maximise the anti-tumour potential of Bt BEVs.

### Intravenously delivered Bt BEVs suppress primary tumour growth in a dose-dependent manner

We next evaluated the therapeutic efficacy of Bt BEVs in primary tumour models. Mice bearing established B16F10 melanoma tumours were treated with increasing doses of BEVs administered IV at three-day intervals following tumour palpation (Fig. 2A). IV administration of Bt BEVs resulted in a clear, dose-dependent inhibition of tumour growth, with the highest dose tested (1×10^10^ BEVs) producing the most pronounced effect (*P* = 0.0185) (Fig. 2B). Tumour growth curves revealed a consistent reduction in tumour progression across the treatment period, and endpoint tumour volumes across multiple independent experiments were significantly decreased in the high-dose group relative to controls (*P* < 0.0001) (Fig. 2C).

**Figure 2.**
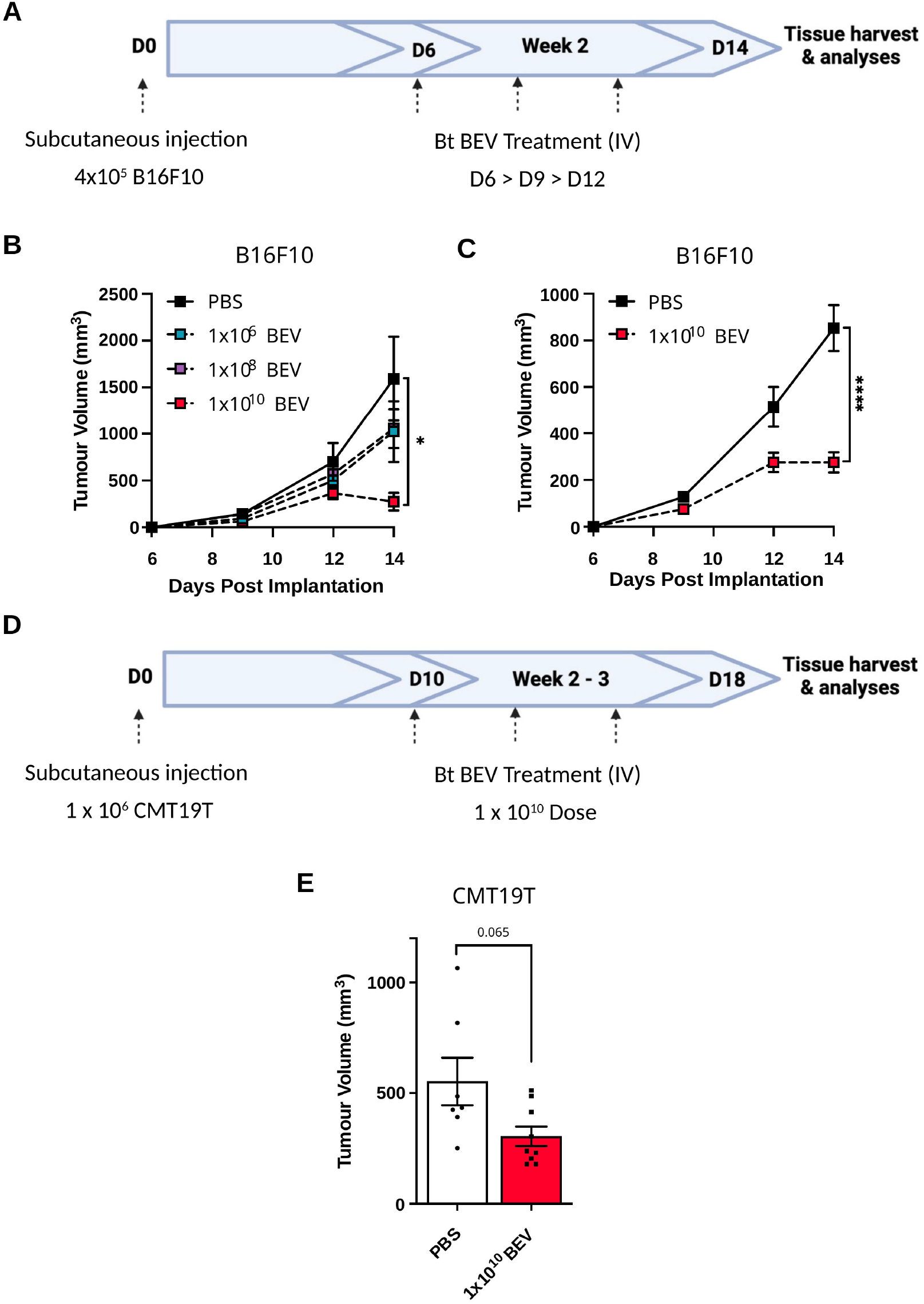
IV Bt BEVs suppress primary tumour growth in a dose-dependent manner. (A) Experimental outline of the B16F10 primary melanoma dose-response study. Mice were inoculated subcutaneously with 4 × 10^5^ B16F10 cells on day 0 and treated IV with Bt BEVs at 3-day intervals from the onset of palpable tumour (days 6, 9, and 12), followed by tissue harvest and analysis on day 14. (B) Mean ± SEM B16F10 tumour growth over time following IV administration of the indicated BEV doses (n = 5). Endpoint dose-response comparisons were analysed by ordinary one-way ANOVA with Dunnett’s multiple-comparisons test against the PBS control. Adjusted *P* values: 1 × 10^6^ BEVs, *P* = 0.4286; 1 × 10^8^ BEVs, *P* = 0.4706; 1 × 10^10^ BEVs, *P* = 0.0185. (C) Mean ± SEM B16F10 tumour growth over time following IV administration of a defined 1 × 10^10^ BEV dose (n = 24, pooled across three independent experiments). Endpoint tumour volume was analysed by Welch’s t-test, P < 0.0001. (D) Experimental outline of the CMT19T primary tumour study. Mice were inoculated subcutaneously with 1 × 10^6^ CMT19T cells on day 0 and treated IV with 1 × 10^10^ Bt BEVs from the onset of palpable tumour, followed by tissue harvest and analysis on day 18. (E) Mean ± SEM endpoint CMT19T tumour volume following IV administration of 1 × 10^10^ Bt BEVs (n = 7–9). Endpoint tumour volume was analysed by Welch’s t-test, *P* = 0.0653. \**P* < 0.05, \*\*\*\**P* < 0.0001.

To assess whether this effect extended beyond a single tumour model, we performed parallel experiments in the CMT19T lung carcinoma model using the same dosing regimen. While the magnitude of response did not reach statistical significance (*P* = 0.0653), BEV-treated animals exhibited the same trend towards reduced tumour burden compared with controls (Fig. 2D-E), suggesting that Bt BEV-mediated anti-tumour activity may extend beyond melanoma. Bt BEV treatment was not associated with overt body-weight loss, and representative H&E-stained sections of major organs showed preserved tissue architecture without obvious inflammatory or necrotic changes (Supplementary Fig. 1). Together, these data demonstrate dose-dependent suppression of primary B16F10 melanoma growth by IV-administered Bt BEVs, with a smaller non-significant reduction in CMT19T lung carcinoma and no obvious histological evidence of treatment-associated tissue damage in the organs examined.

### Bt BEVs inhibit metastatic tumour outgrowth in a preclinical melanoma model

Given the clinical importance of metastatic progression in melanoma, we next investigated whether Bt BEV treatment impacts secondary tumour outgrowth. To model metastatic disease, B16F10 melanoma cells were introduced IV to allow seeding in the lung, followed by initiation of BEV treatment after tumour cell dissemination (Fig. 3A). At experimental endpoint, visual inspection of lung tissue revealed fewer and smaller pulmonary nodules in BEV-treated animals compared to controls. Quantitative analysis showed a reduction in the number of pulmonary lesions (Fig. 3B), approaching statistical significance (*P* = 0.0549), whilst representative H&E-stained lung sections were consistent with reduced lesion size in BEV-treated animals (*P* = 0.011) (Fig. 3C). These findings indicate that BEV treatment limits metastatic tumour outgrowth, even when administered after initial tumour cell seeding. These results extend the anti-tumour activity of Bt BEVs beyond primary tumour growth and suggest that BEV treatment impacts multiple stages of tumour progression.

**Figure 3.**
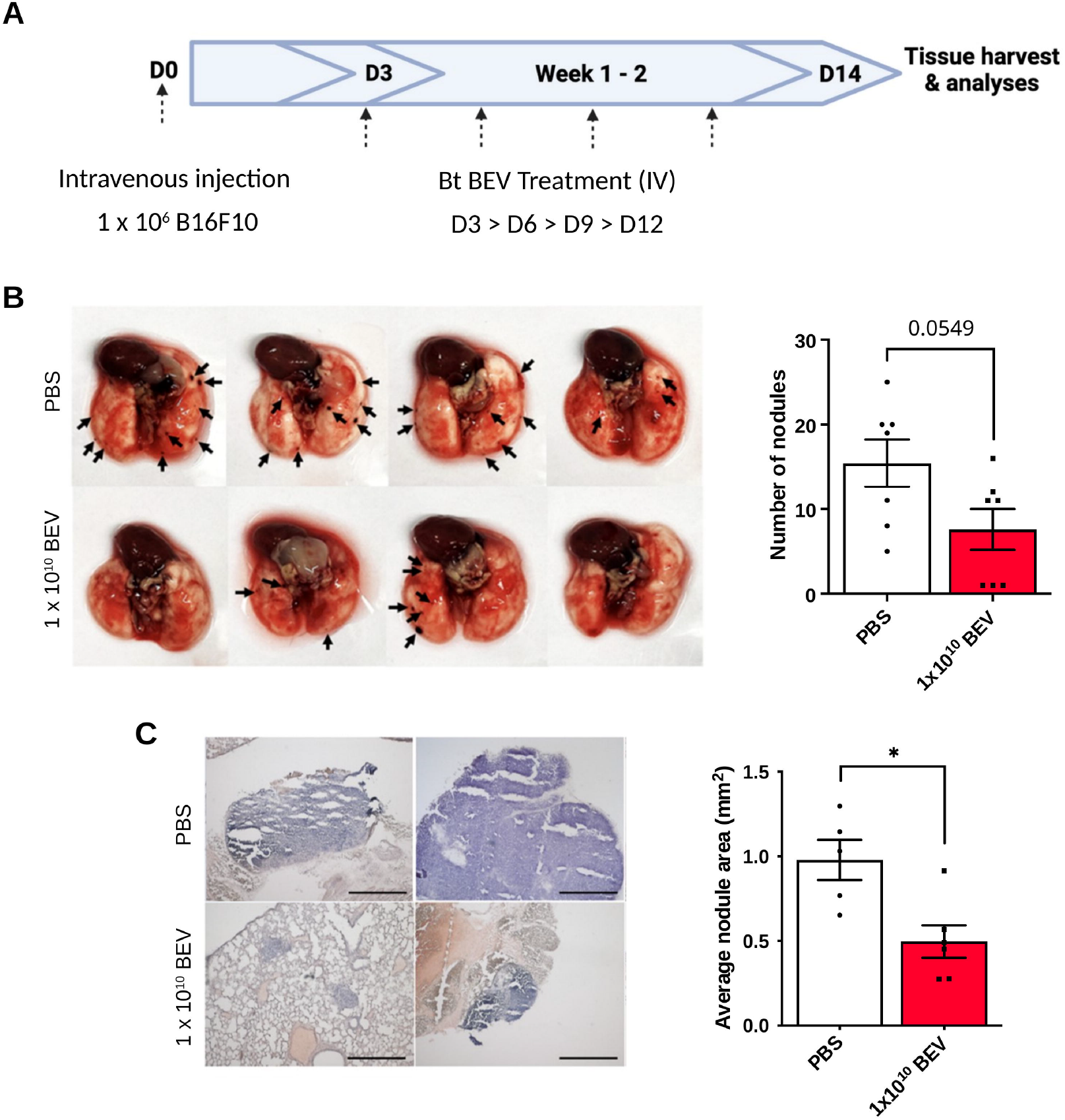
Bt BEVs reduce pulmonary lesion burden in a melanoma experimental metastasis model. (A) Experimental outline of the B16F10 experimental metastasis study. Mice were injected IV with 1 × 10^6^ B16F10 cells on day 0 and allowed to seed in the lungs for 3 days before initiation of IV Bt BEV therapy (1 × 10^10^ BEVs) on days 3, 6, 9, and 12, followed by tissue harvest and analysis on day 14. (B) Representative lung images (left) and quantification of the number of visible pulmonary surface lesions (right) following vehicle control or Bt BEV treatment. Bars represent mean ± SEM, n = 7. Two-tailed unpaired t-test, *P* = 0.0549. (C) Representative H&E-stained lung sections acquired at 4× objective (left) and quantification of average pulmonary lesion area (right) in animals treated with vehicle control or 1 × 10^10^ Bt BEVs. Scale bars, 200 µm. Lesion area was quantified in ImageJ. Bars represent mean ± SEM, n = 5–6. Two-tailed unpaired t-test, *P* = 0.0110. \**P* < 0.05.

### Bt BEVs activate innate immune signalling and localise to tumour tissue following systemic administration

To explore potential mechanisms underlying the observed anti-tumour effects, we first assessed whether Bt BEVs directly impact tumour cell viability. Co-culture of B16F10 cells *in vitro* with increasing concentrations of BEVs did not reduce tumour cell viability (*P* = 0.342) (Fig. 4A), consistent with the anti-tumour effects observed *in vivo* not being due to direct cytotoxicity. We next evaluated the capacity of Bt BEVs to stimulate innate immune signalling pathways. BEV co-culture with THP-1-Blue cells induced robust activation of NF-κB signalling in a dose-dependent manner (Fig. 4B). Consistent with this, Bt BEVs activated both TLR2 and TLR4 in HEK-Blue reporter cells, with a stronger response through TLR2 (Fig. 4C–D), indicating engagement of canonical innate immune sensing pathways. Finally, to determine whether systemically administered BEVs can access tumour tissue, we performed biodistribution studies using NanoLuc-expressing Bt BEVs (BEV-NLuc). Following IV administration, BEV-NLuc signal was detected in multiple organs, including within tumour tissue (Fig. 4E-G), consistent with tumour localisation *in vivo*. Together, these findings demonstrate that Bt BEVs are potent activators of innate immune signalling and can localise to tumour sites following systemic administration, providing a plausible basis for their observed anti-tumour activity.

**Figure 4.**
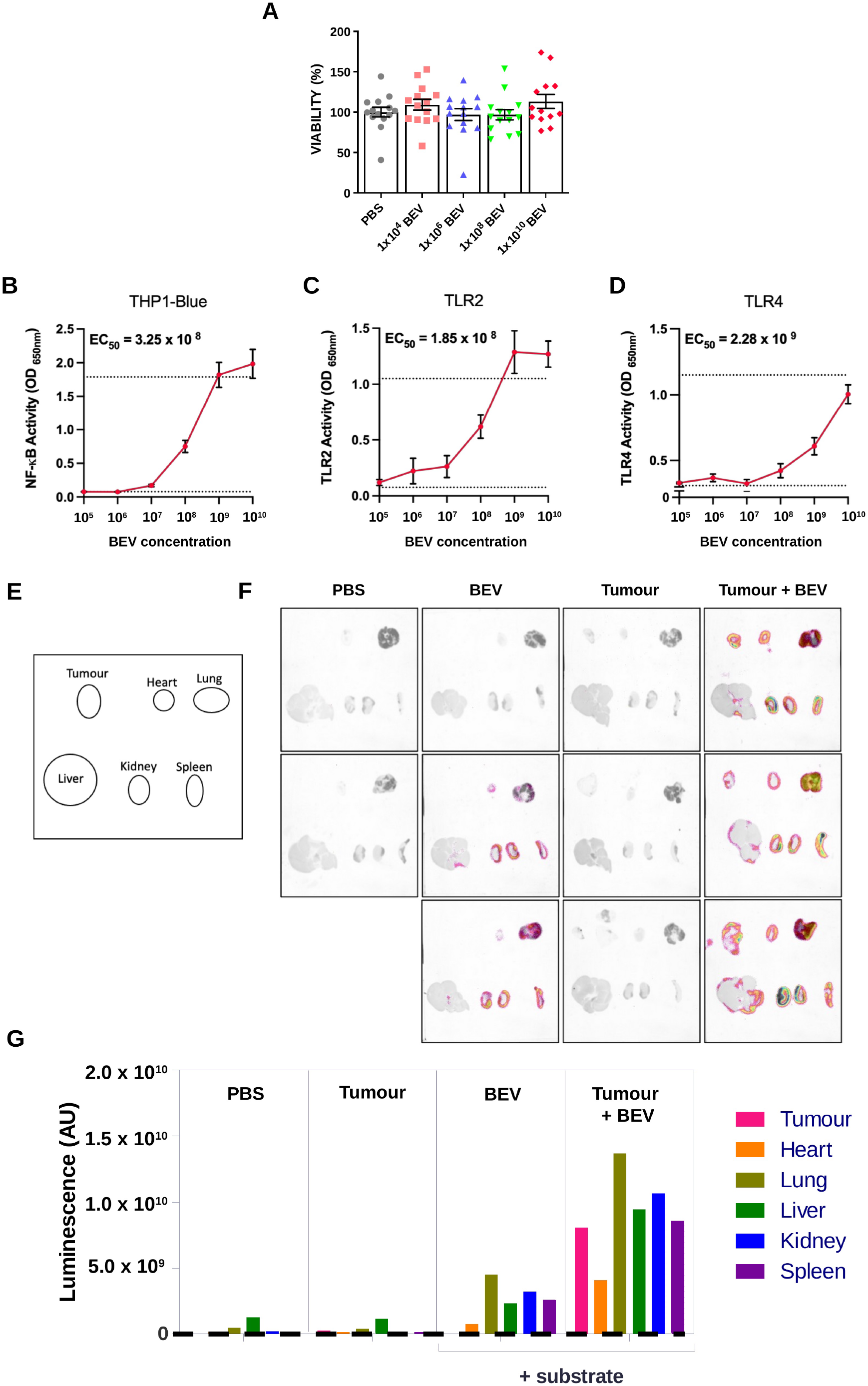
Bt BEVs activate innate immune signalling and localise to tumour tissue. (A) Relative B16F10 tumour-cell viability following *in vitro* treatment with the indicated BEV doses. Viability was quantified by alamarBlue fluorescence, n = 15, N = 3. Statistical significance was calculated by ordinary one-way ANOVA; no significant difference was detected among treatment groups, *P* = 0.3425. (B) Quantification of NF-κB activity in THP1-Blue monocytes in response to escalating BEV doses. Activity was measured by QUANTI-Blue detection of the SEAP reporter, n = 5. (C) Quantification of TLR2 activity in HEK-Blue hTLR2 cells in response to escalating BEV doses, n = 3. (D) Quantification of TLR4 activity in HEK-Blue hTLR4 cells in response to escalating BEV doses, n = 3. (E) Schematic showing the *ex vivo* organ layout used for biodistribution imaging. (F) Representative *ex vivo* bioluminescence images of organs from non-tumour-bearing or day 10 B16F10 tumour-bearing mice following IV administration of vehicle control or 1 × 10^10^ NanoLuc-tagged BEVs (BEV-NLuc). (G) Quantification of absolute bioluminescent signal across indicated organs following vehicle or BEV-NLuc administration, n = 3. For biodistribution studies, organs were harvested 3 h after IV dosing and imaged following incubation in furimazine substrate.

## Discussion

Our data show that BEVs derived from the human intestinal commensal *Bacteroides thetaiotaomicron* exert anti-tumour activity *in vivo* that is strongly influenced by delivery route. In the preclinical melanoma model examined here, IV administration, but not IP delivery, produced marked suppression of primary B16F10 melanoma growth and reduced pulmonary lesion burden in an experimental lung colonisation model. In the absence of a direct cytotoxic effect on tumour cells *in vitro*, these findings are consistent with BEVs eliciting a host immune-mediated mechanism of action. Prior anti-cancer BEV studies have utilised non-commensal or engineered systems, making the present study distinct in its focus on native vesicles from a human commensal bacterium (*12, 24*).

A key conclusion of this study is that the delivery route is a major determinant of Bt BEV anti-tumour efficacy. This finding aligns with the broader BEV literature showing that biological activity depends not only on vesicle dose or source organism, but also on the route by which vesicles encounter host tissues and immune compartments. In cancer models, *Kim et al*.(*12*) showed that IV-administered BEVs from attenuated pathogens accumulate in tumours and induce anti-tumour immune responses. By contrast, engineered anti-EGFR BEVs derived from non-pathogenic *E. coli* BL21(DE3) have shown anti-tumour activity after IP administration in a 4T1 breast cancer model(*24*), whilst bioengineered commensal-derived Bt BEVs delivered orally or intranasally can elicit local and systemic immune responses in vaccine settings(*20, 25*). Recent preclinical cancer studies have further shown that orally delivered commensal or engineered BEVs can also elicit anti-tumour responses through distinct mucosal or vaccine-like mechanisms(*26, 27*). Thus, the optimal route for BEV-based therapeutics is likely context dependent, reflecting differences in vesicle source, engineering strategy, cargo, tissue exposure, and intended immune mechanism. In our model, IV delivery was superior to IP delivery, but the pharmacokinetic and cellular basis of this route dependence remains to be defined.

Although the present study does not identify the *in vivo* mechanisms, the data are supportive of an innate immune-modulatory model of BEV activity. Bt BEVs activate NF-κB and TLR reporter systems *in vitro*, and NanoLuc-labelled BEVs localise to tumour tissue after IV administration. This is consistent with prior work showing that Bt BEVs modulate dendritic-cell responses in healthy states(*8*), and with evidence that Bt BEVs may reprogramme innate immune responses in THP-1 Blue monocytes through TLR2-associated changes in DNA methylation(*16*). Importantly, the *Durant et al*.(*8*) study also showed that Bt BEV-induced dendritic-cell responses differ between healthy and IBD-derived cells, indicating that host inflammatory context can shape responsiveness to Bt BEVs. Tumour-bearing tissues represent a distinct disease context from intestinal inflammation, and the inflammatory composition of the tumour microenvironment may therefore shape which innate immune populations respond to Bt BEV exposure. In melanoma models, agonist strategies engaging TLR1/2(*28*) and TLR2(*29*) have been shown to enhance anti-tumour immunity through mechanisms involving dendritic cell activation, co-stimulation, IFN-γ-associated CD8 T-cell responses, and improved checkpoint blockade activity. Together, these data support a biologically plausible innate immune-mediated model of Bt BEV activity in this tumour setting, with future work required to define the responsible effector pathways.

This study has some notable limitations. The strongest efficacy data were generated in B16F10 melanoma, with a smaller, non-significant reduction observed in CMT19T lung carcinoma, indicating that Bt BEV activity may be tumour-context dependent. In addition, our data do not identify specific BEV component(s), receptor pathways, or immune cell circuits responsible for the *in vivo* phenotype. Nonetheless, this work provides proof-of-principle that native commensal Bt BEVs can suppress tumour progression *in vivo* and identifies the delivery route as a critical determinant of efficacy. More broadly, these findings support the concept that microbiome-derived vesicles can mediate systemic host–tumour effects beyond the intestinal niche, offering a tractable cell-free route for exploiting host– microbe immune interactions without requiring live bacterial administration.

## Acknowledgements

This work was supported by funding from: BigC (grant number 18-15R); SDR acknowledges the support of the Biotechnology and Biological Sciences Research Council (BBSRC). SRC, SDR, RJ and RS gratefully acknowledge the support of the Biotechnology and Biological Sciences Research Council (BBSRC); this research was funded by the BBSRC Institute Strategic Programme Grants Gut Microbes and Health (BB/R012490/1) and its constituent projects (BBS/E/F/000PR10353, BBS/E/F/000PR10355, and BBS/E/F/000PR10356), and Food Microbiome and Health BB/X011054/1 and its constituent project(s) BBS/E/QU/230001B, BBS/E/QU/230001C.

## Author contributions

Conceptualisation: CAP, LJH, SRC SDR; Formal analyses: CAP, EJJ, NI. Investigation: CAP, EJJ, NI, AN, AMM, LM, RS; Resources: LJH, SRC, SDR; Review and editing: CAP, RS, SRC, SDR; Visualisation: CAP, EJJ, NI, RS; Supervision: LJH, SRC, SDR; Funding acquisition: LJH, SRC, SDR.

