## Supplementary figures and images for "Systemically delivered *Bacteroides thetaiotaomicron*-derived bacterial extracellular vesicles inhibit primary and metastatic melanoma growth"

### Supplementary Figure 1

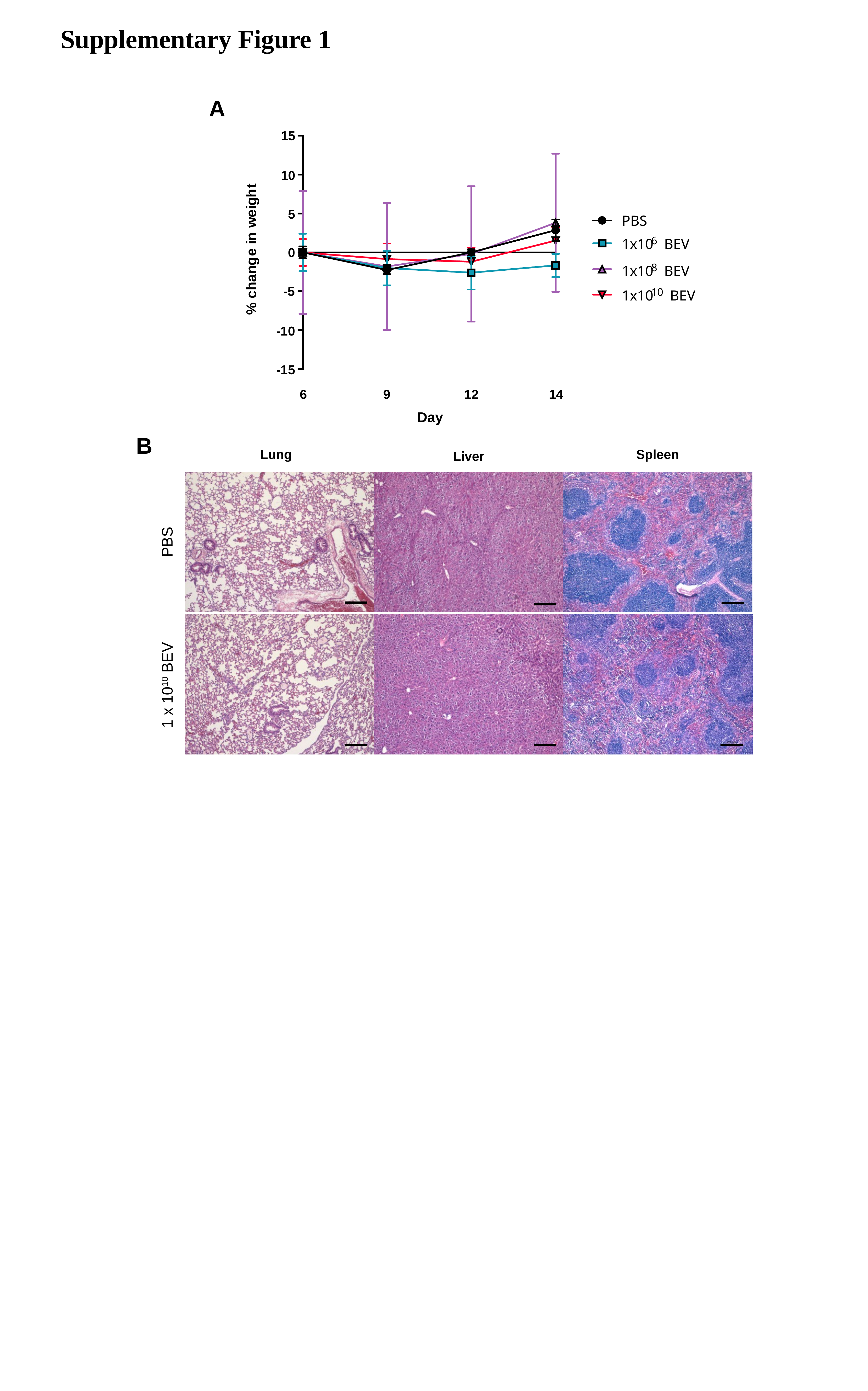
